## Supplemental materials Figs. S1-S4 for "An Engineered Nanocomplex with Photodynamic and Photothermal Synergistic Properties for Cancer Treatment"

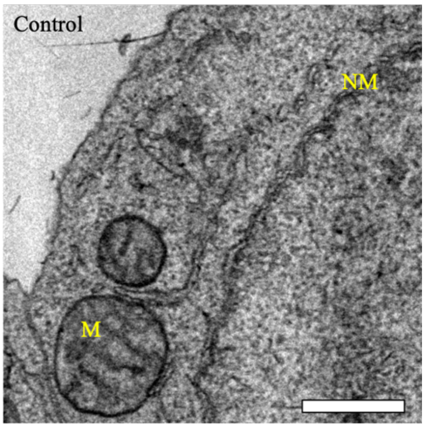


**Figure S1 | SH-SY5Y untreated TEM images (A)** Electron micrograph of SH-SY5Y untreated cells. The image shows mitochondria (M) that do not contain AuNP-mTHPC. The plasma membrane and the nuclear membrane (NM) are well shaped. Scale bar = 0.5 µm.

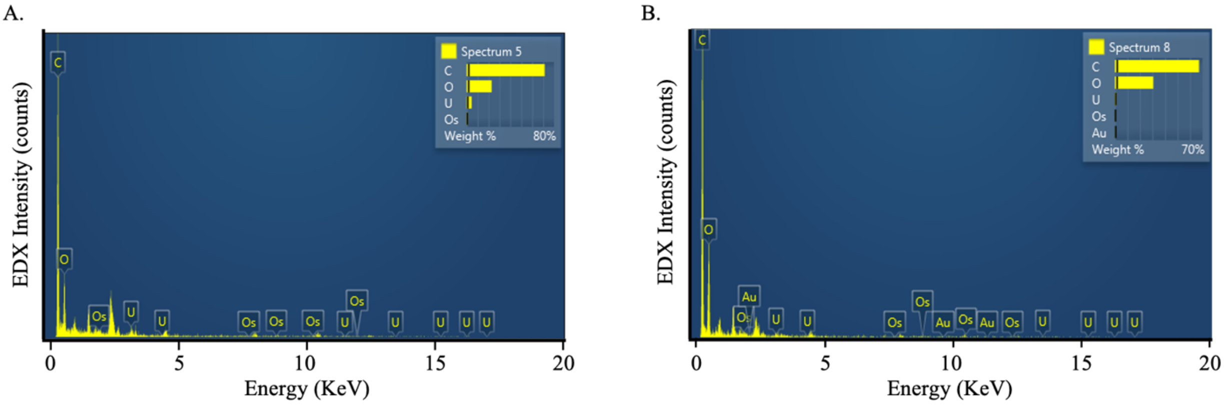

**Figure S2 | EDX analysis of SH-SY5Y cells.** Energy disperse spectroscopic spectra **(A)** Untreated cells (B) Cells treated with 1.2 µM AuNP-mTHPC. The element weight percent of C, O, Os, Au, and U is displayed.

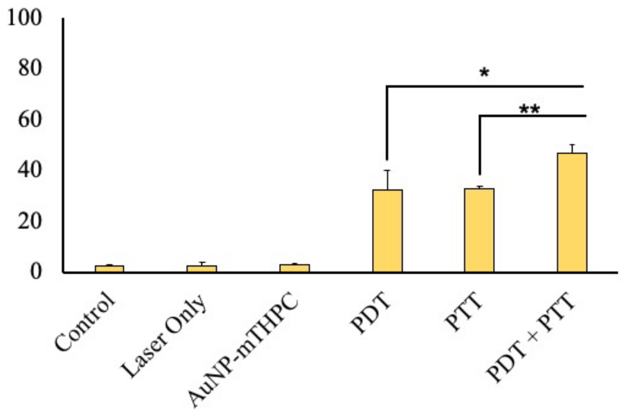


**Figure S3 |** Cell death induced by laser irradiation after 24 h incubation with AuNP−mTHPC complex. **(A)** Cell viability of SH-SY5Y cells with 1.2 µM AuNP-mTHPC and illuminated under 650 nm laser (PDT) at 6 mW/cm$^2$ for 4 min or illuminated under 532 nm laser (PTT) at 15 mW/cm$^2$ for 4 min. The combination of PDT PTT was conducted using 2 min of each laser. Average quantification of three experiments is presented in the plots (mean ± STDEV). There were significant differences in the relative levels control and PDT and PTT in the experiments. **P < 0.01, *P < 0.05.

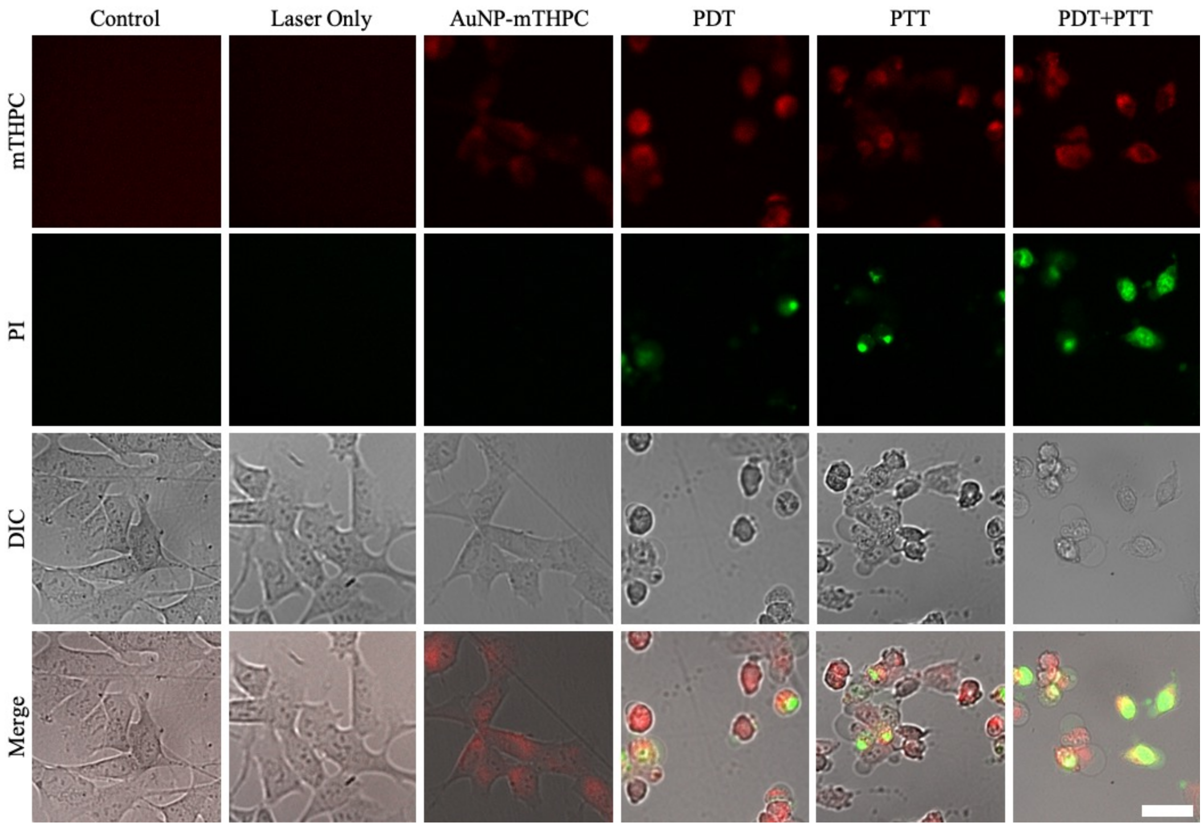


**Figure S4 |** Microscopy Images of SH-SY5Y cells after PDT/PTT treatment **A)** Confocal microscopy of 1.2 μM AuNP-mTHPC (red) and PI (green) labeled SH-SY5Y cells after laser irradiation under 650 nm laser (PDT) at 6 $mW/cm^2$ for 4 min or/and illuminated under 532 nm laser (PTT) at 15 $mW/cm^2$ for 4 min. Scale bar = 50 μm.
